# Enzyme-Linked Cycloaddition Assay (ELCA) for rapid, ultra-sensitive monitoring of secreted sialoglycoproteins

**DOI:** 10.64898/2026.06.15.732414

**Authors:** Jon Lundstrøm, Jingmei Yang, Daniel Bojar

## Abstract

Glycosylation of proteins is central to cell signaling, immune function, and pathogen interactions, yet existing methods for monitoring glycan changes require specialized instrumentation and primarily report on membrane-anchored, rather than secreted, glycoproteins, with a slow turnover. Here, we present the Enzyme-Linked Cycloaddition Assay (ELCA), a click chemistry-based platform for ultra-sensitive detection and semi-quantitative analysis of secreted sialoglycoproteins. By metabolically incorporating an azide-modified sialic acid into newly synthesized glycoproteins and capturing labeled material via strain-promoted cycloaddition, ELCA quantifies aggregate sialylation using a microplate reader-compatible, ELISA-like workflow. We demonstrate that the secreted glycoproteome responds rapidly to pharmacological perturbation, with changes detectable within hours. Benchmarking against common glycosylation inhibitors and profiling cytokine-driven macrophage polarization further establishes ELCA’s sensitivity and temporal resolution. Compatible with serum-containing conditions and requiring no specialized instrumentation, ELCA provides a broadly accessible tool for rapid, cost-effective monitoring of secreted glycoprotein dynamics.

## Introduction

Glycans modify the vast majority of proteins that traverse the secretory pathway, including both membrane-anchored and secreted proteins^1^. As complex carbohydrate structures exposed to the extracellular environment, glycans play central roles across cell–cell and cell–pathogen interactions^2,3^, tumor immune evasion^4^, microbiome development^5^, and further regulate the stability, structure, and function of the proteins they modify^6^. Overall, glycosylation is involved in diverse biological processes and is increasingly recognized as an important target in both diagnostic and therapeutic applications^7,8^.

Glycans are enzymatically synthesized in a non-templated manner, where the availability and flux of protein substrates, glycosyltransferase enzymes, and monosaccharide donors collectively determine the final structures. Investigating the cellular glycome remains inherently challenging, due to extreme structural diversity arising from the assembly of chemically highly similar monosaccharide building blocks^9^.

Comprehensive structural characterization using mass spectrometry-based approaches is powerful but typically expensive, time-consuming, and dependent on access to specialized instrumentation and expert domain knowledge. Further, representative insights into global motif abundance, such as sialylation, requires the acquisition of multiple types of glycomics data (*N*-, *O*-glycans, glycosphingolipids)^10^. Alternatively, glycan profiling using recognition by specific glycan-binding proteins (lectins) as a proxy serves as a viable alternative, as lectin binding patterns can be directly correlated with the glycan motifs present in a sample and often aggregate information from several glycan classes^11^, yet suffers from variable specificity of lectins for binding their motif in certain sequence contexts.

Secreted glycoproteins, including cytokines, hormones, growth factors, enzymes, etc., are particularly attractive as a source of material for monitoring cellular activity, allowing for continuous measurements with little to no sample preparation requirements as well as rapid response to changes in the cellular state^12^. In contrast, the glycan profile of membrane-anchored proteins responds more slowly to perturbations, since pre-existing surface proteins must first be turned over before changes are observed^13^. However, the analysis of secreted glycoproteins presents several challenges, including relatively low abundances and potential interference from serum-derived glycoproteins under standard culture conditions, which is why most methods investigate glycosylation levels from the cell lysate (e.g., mass spectrometry) or the membrane itself (e.g., flow cytometry).

Previous studies have developed mass spectrometry-based protocols for quantifying secreted (glyco)proteins under standard culture conditions, combining bioorthogonal metabolic labeling with specific enrichment and TMT-based multiplexing to boost signal sensitivity^14,15^. While such approaches enable detailed protein-level quantification, they come with the trade-off of being cost- and labor-intensive, and may not be readily accessible to many laboratories. Although site-specific glycosylation can be inferred with such methods, the reported results are typically limited to aggregate *N*-glycosylation, as quantification relies on the comparison of peptide signals before and after enzyme-mediated deglycosylation, without providing information on glycan composition. Importantly, it was demonstrated that click chemistry-based enrichment of secreted glycoproteins alone is insufficient for comprehensive analysis, as non-specifically retained serum-derived proteins can still dominate the signal^15^.

Several ELISA-derived methods, such as the Enzyme-Linked Lectin Assay (ELLA)^16^ and Enzyme-Linked Lectin Binding Assay (ELBA)^17^, offer more cost-effective and accessible approaches for quantifying soluble glycoproteins. However, these methods require serum-free culture conditions, as glycan motifs indistinguishable from those of the sample of interest are present in highly abundant serum-derived proteins.

Here, we develop and benchmark a method to detect and quantify rapid changes in the secreted glycoproteome. While higher-resolution approaches (e.g., LC-MS/MS-based glyomics and/or glycoproteomics) can provide detailed structural information, they require substantially more complex experimental setups, as well as time and resource investments. In contrast, our herein developed Enzyme-Linked Cycloaddition Assay (ELCA) enables monitoring global changes in secreted glycoproteins (total sialylation, i.e., change in total glycoprotein and/or change in glycosylation) using a simple and cost-effective workflow using click chemistry. The assay is comparable to a standard ELISA setup and requires only access to a microplate reader, thereby facilitating higher-throughput analyses across replicates, time points, and treatment conditions than existing methods and opening up sensitive glycan analyses to all molecular biology laboratories. Importantly, the per-sample cost is as low as 1-2 EUR, making it orders of magnitude less expensive than mass spectrometry-based analyses.

Using ELCA, we demonstrate here that changes in the secreted glycome are detectable at substantially earlier time points compared to analyses of cell surface–associated glycoproteins. We evaluate the effects of common glycosylation inhibitors and observe rapid impacts on secreted glycoproteins, while conventional cell surface analysis by flow cytometry requires considerably longer treatment durations to reveal comparable changes, providing caveats and guidelines to current usage of such inhibitors in scientific studies. Finally, we characterize the secreted glycoprotein response of THP-1-derived macrophages following cytokine-mediated activation, yielding insights into the early response of glycoprotein secretion. Overall, we envision ELCA as a general platform for assessing the impact of treatments on glycosylation in a rapid, cost-effective, and sensitive manner.

## Results

### ELCA enables semi-quantitative analysis of secreted glycoproteins under standard culture conditions

Sialylation, capping glycans with one or multiple sialic acids, is one of the most prominent and relevant glycan features in mammals and thus highly studied for many biological questions. It has been shown to control the half-life of secreted proteins^18^, as well as mediate interactions with immune system proteins such as Siglecs^19^ and, being tied to the same biosynthetic machinery as membrane-associated glycoproteins, is a proxy for general mature cellular glycosylation^19^. We thus aimed to develop a method that would allow any researcher to quantify time-resolved changes in sialylation in a sensitive and cost-effective manner that did not require specialized instrumentation. To achieve this aim, such a method had to overcome several hurdles, such as the presence of similar, or even identical, glycans on FBS-derived glycoproteins in cell culture, or the slow time-scales of plasma membrane turnover. Additionally, we caution that, even using mass spectrometry, researchers usually focus on only one type of glycosylation, such as *N*- or *O*-glycomics, and do not necessarily observe changes in the total glycome of a cell.

Taken all these considerations together, we decided to use metabolic labeling to introduce azide groups into sialic acids (Neu5Az, instead of Neu5Ac) that we could then use for bioorthogonal click chemistry to introduce biotin tags that would allow for sensitive detection of secreted sialoglyoproteins, without interference of non-azide-containing serum glycoproteins, in a method we termed Enzyme-Linked Cycloaddition Assay (ELCA, Fig. 1A). We note that, thanks to the incorporation of Neu5Az into all glycan classes^20^, ELCA thus reported on global sialoglycoprotein levels, combining signals from both *N*- and *O*-glycans, as well as all sialic acid linkages.

**Figure 1.**
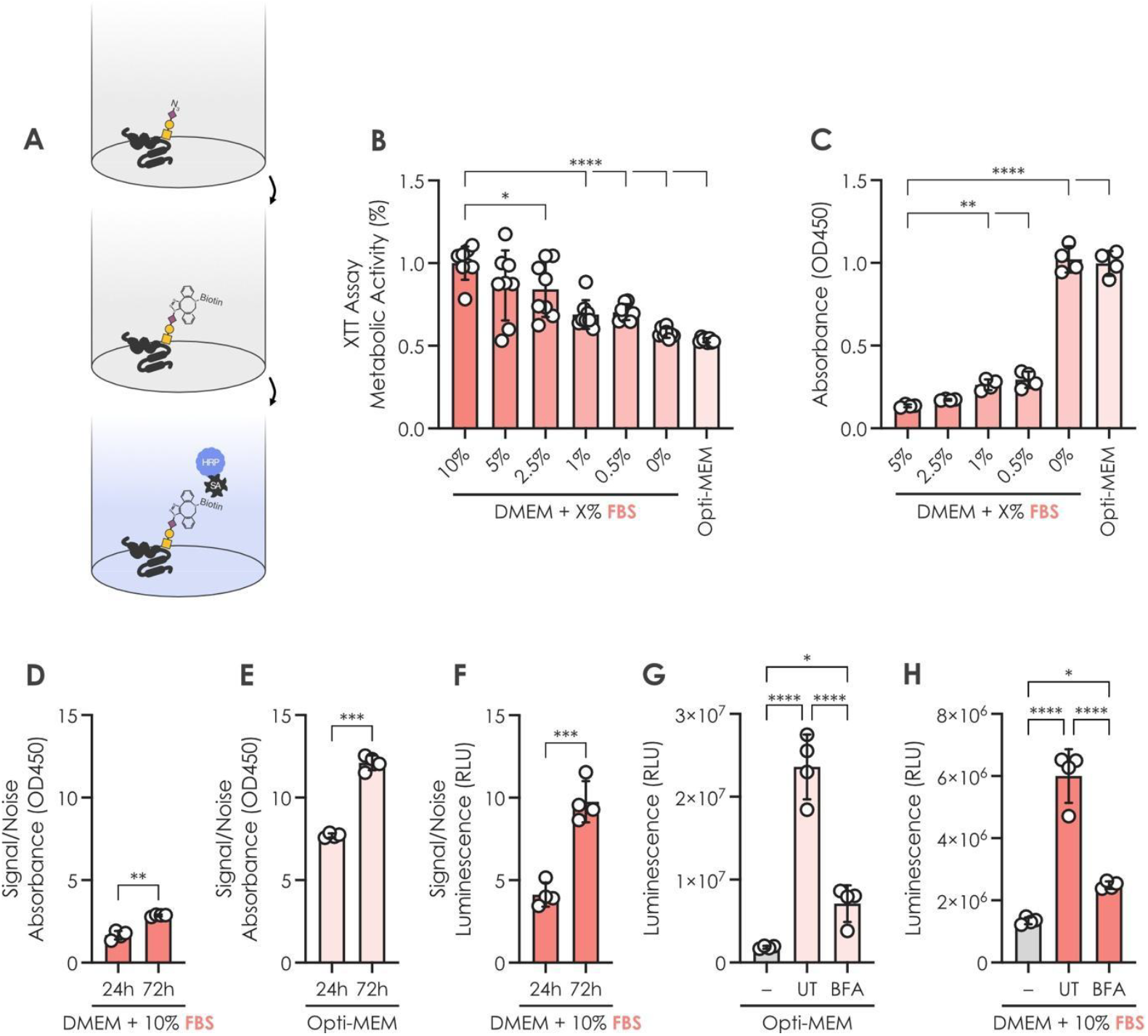
Quantifying secreted glycoproteins under standard culture conditions. **A)** Schematic of the ELCA workflow. Supernatant samples containing azide-modified glycoproteins are adsorbed onto microwell plates and subsequently labeled via strain-promoted cycloaddition. Detection and quantification are performed using HRP. **B)** XTT assay quantifying cellular metabolic activity under culture conditions with varying serum concentrations. (n = 8). **C)** ELCA analysis of supernatants from cells cultured under varying serum concentrations using standard colorimetric detection (n = 4). **D-E)** Signal-to-noise ratio (S/N) of ELCA signal comparing standard culture conditions (DMEM + 10% serum; D) and serum-free Opti-MEM (E) at 24 h and 72 h time points, measured by standard colorimetric detection (n = 4). **F)** S/N of ELCA signal from cells cultured under standard conditions at 24 h and 72 h timepoints, measured using high-sensitivity luminescent detection (n = 4). **G-H)** ELCA signal following treatment with 3 µM Brefeldin A (BFA) for 24 h under serum-free (G) or standard culture conditions (H), measured using high-sensitivity luminescent detection (n = 4). – (grey bars): blank control. All assays were conducted in A375 cells. Statistical significance was assessed using a two-tailed Welch’s t-test (two conditions) or one-way ANOVA with Dunnett’s multiple comparisons test (>2 conditions, 1 variable) *p < 0.05, **p < 0.01,

With the goal of quantifying secreted glycoproteins under standard culture conditions, including the presence of serum, we set out to optimize the ELCA workflow for maximal sensitivity. We found that careful titration of horseradish peroxidase (HRP), as well as the choice of blocking reagent and protocol, was critical for minimizing non-specific background signal and achieving an optimal signal-to-noise ratio (S/N; Fig. S1A-B). We note that these parameters may require validation and further optimization when switching reagent lots or establishing the assay in a new laboratory.

While serum starvation would reduce interference from highly abundant background proteins, we confirm that cellular metabolic activity is significantly affected under these conditions (Fig. 1B), which risks introducing artificial changes into the secreted glycoproteome. Moreover, while short-term serum starvation may be tolerated, the feasibility of time-course experiments to monitor additional perturbations, e.g., small-molecule treatments, would be limited.

Using standard colorimetric detection, we observed that the sensitivity of this first version of ELCA was significantly reduced in the presence of even small amounts of exogenous serum-derived proteins (Fig. 1C). This effect was likely due to competition or crowding, as the assay relies on coating the microwell plate with the entire, complex sample. Extending the culture duration from 24 h to 72 h provided small improvements, yet the overall signal-to-noise ratio and dynamic range remained low under serum-containing conditions, compared to serum-free conditions (Fig. 1D-E).

To address this limitation, we evaluated alternative detection strategies and found that chemiluminescent substrates designed for ultra-sensitive detection substantially improved signal detection in both serum-containing and serum-free conditions (Fig. 1F, Fig. S1C). With the example of Brefeldin A (BFA) treatment, effectively blocking secretion, we then observed a strong signal in control conditions that was reduced to near-background levels upon treatment, both in serum-containing and serum-free supernatants (Fig. 1G-H).

We confirmed that the detected signal is specific to sialylated glycoproteins upon metabolic labelling with Ac_4_ManNAz. Specifically, (i) no non-specific labeling by DBCO occurred in the absence of metabolic labeling, (ii) no measurable signal was observed from medium containing Ac_4_ManNAz but no cells, and (iii) the signal was significantly reduced upon treating supernatants with a broad-specificity sialidase prior to analysis (Fig. S1D). We also evaluated metabolic labeling using Ac_4_GalNAz in a proof-of-concept experiment. Ac_4_GalNAz primarily labels GalNAc-containing glycans (e.g., *O*-GalNAc), but can also contribute to labeling of GlcNAc-containing structures (including *O*-GlcNAc and *N*-glycans) through GALE-mediated epimerization to GlcNAz^21,22^. Consistent with this, we detected a strong ELCA signal in supernatants from Ac_4_GalNAz-treated cells that was insensitive to sialidase treatment (Fig. S1D).

To further expand the potential applications of the method, we finally explored the quantification of a specific secreted glycoprotein, using GM-CSF, a heavily glycosylated cytokine known to be induced in A375 cells upon PMA treatment^23^, as an example. Using the standard ELCA workflow, we first observed a general decrease in total secreted glycoproteins upon PMA treatment (Fig. S1E, in grey). Interestingly, when applying an ELISA-inspired sandwich setup, using a monoclonal antibody for capture followed by click-based detection of azide-labeled glycans, we detected a concentration-dependent increase in glycosylated GM-CSF (Fig. S1E, in red), demonstrating that our method can be adapted to monitor specific glycoproteins, in addition to global secretion profiles.

### ELCA detects changes in the secreted glycoproteome with high temporal resolution

The cell surface glycome reflects the state of a given cell. For example, dysregulated metabolic activity, such as during stress responses or cancer, has been shown to alter the composition of glycans presented to the extracellular environment. These changes can confer advantages to cancer cells by modulating interactions with immune cells, while also serving as potential biomarkers and therapeutic targets^24^.

Because measurable changes in the cell surface glycome depend on the turnover of existing glycoproteins, we hypothesized that the abundance or rate of glycoprotein secretion could provide a more sensitive readout for early detection of cellular perturbations. Consistent with this, brief treatment with cycloheximide (CHX) to inhibit protein translation did not result in detectable changes in the surface glycome within 24 h, as assessed by lectin staining (longer treatments were not feasible due to reduced cell viability) (Fig. 2A-B). In contrast, analyzing the supernatant using ELCA revealed a clear reduction in newly secreted glycoproteins under the same conditions (Fig. 2C).

**Figure 2.**
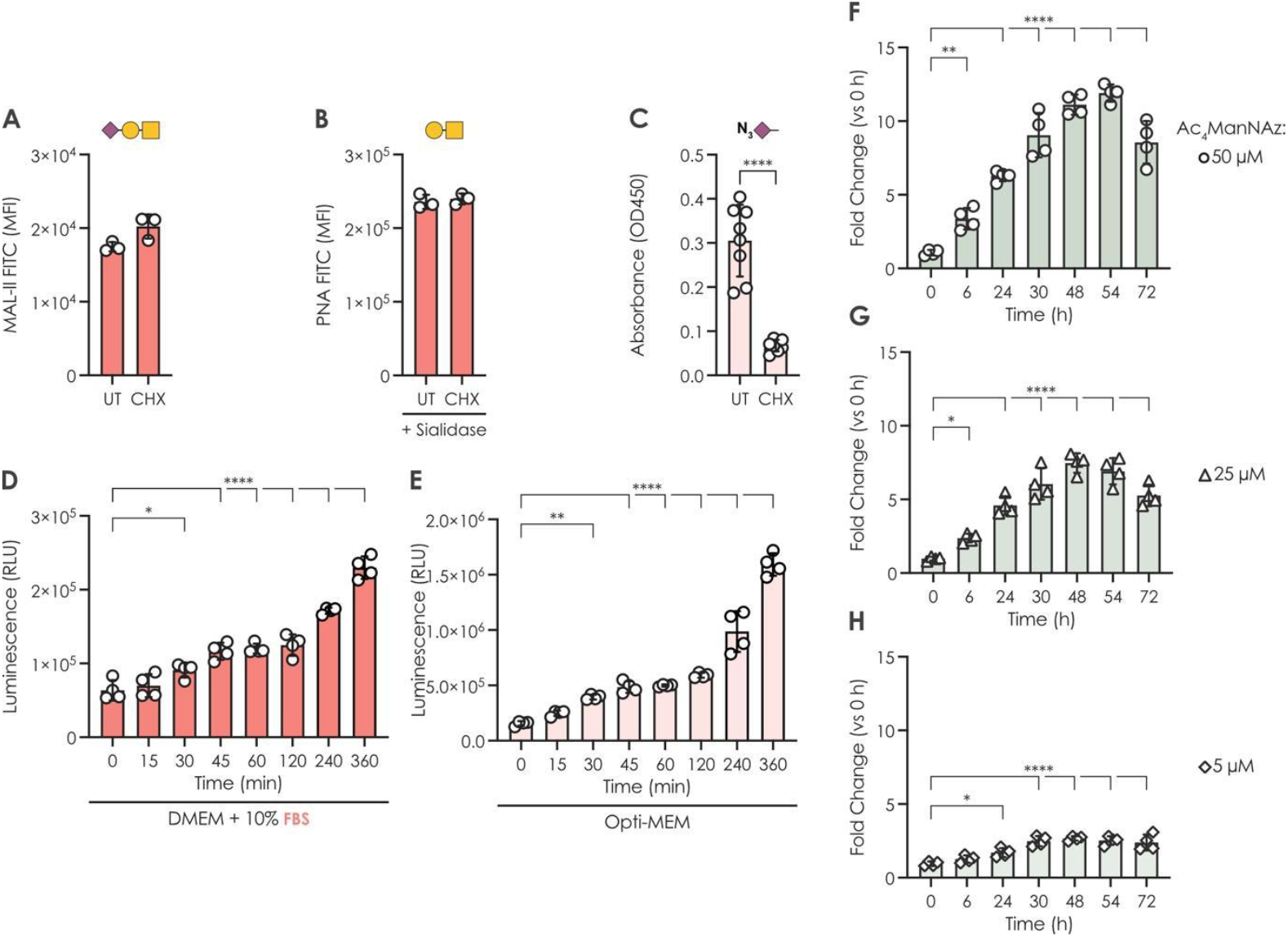
ELCA detects rapid changes in the secreted glycoproteome. **A-B)** Lectin-based flow cytometry analysis of cell surface glycoproteins using MAL-II (A; Neu5Acα2-3Galβ1-3GalNAc) and PNA (B; Galβ1-3GalNAc) lectin staining following inhibition of translation with 25 µM cycloheximide (CHX) for 24 h showed no significant changes (n = 3). **C)** Under the same treatment conditions, a clear reduction in newly secreted sialoglycoproteins was detected using ELCA. Measurements were performed under serum-free conditions using standard colorimetric detection (n = 8). **D-E)** A375 cells were metabolically labeled with 50 µM Ac_4_ManNAz for 72 h, followed by re-seeding in fresh serum-containing DMEM (D) or serum-free Opti-MEM (E) medium. Supernatant samples were collected at the indicated time-points and assayed with ELCA using high-sensitivity luminescent detection (n = 4). **F-G)** Serum-containing DMEM medium supplemented with Ac_4_ManNAz (F: 50 µM; G: 25 µM; H: 5 µM) was added to A375 cells. At the indicated time points, the cells were then incubated in fresh serum-free Opti-MEM for 1 h, after which supernatants containing proteins secreted during this period were collected and stored at −20 °C for ELCA analysis using high-sensitivity luminescent detection (n = 4). Glycan structures have been drawn with GlycoDraw^25^ and conform with the Symbol Nomenclature For Glycans^26^ (SNFG). All assays were conducted in A375 cells. Statistical significance was assessed using a two-tailed Welch’s t-test (two conditions) or one-way ANOVA with Dunnett’s multiple comparisons test (>2 conditions). *p < 0.05, **p < 0.01, ****p < 0.0001.

To benchmark the sensitivity of ELCA, we next investigated the minimum secretion time required for reliable detection of glycoproteins in the supernatant. Under both standard culture conditions (Fig. 2D) and serum-free conditions (Fig. 2E), a significant and specific signal was detectable after just 30 minutes of incubation with fresh medium, demonstrating the high sensitivity and temporal resolution of ELCA, even in the presence of highly abundant serum-derived proteins.

Finally, leveraging this sensitivity and ability to detect rapid changes, we next applied ELCA to monitor the dynamics of Ac_4_ManNAz metabolic labeling across different concentrations and time points. In A375 cells, we detected the secretion of azide-tagged glycoproteins as early as our first timepoint (six hours) following the addition of Ac_4_ManNAz to the culture medium (Fig. 2F-G). Furthermore, we were able to track labeling efficiency over time and observed time- and concentration-dependent saturation with maximal labelling of newly secreted glycoproteins observed after around two days of incubation (48 - 54 hours; Fig. 2F-H). Interestingly, the observed secretion rate signal decreased again at 72 hours (Fig. 2F-G), highlighting the importance of considering azido-sugar consumption and half-life during longer experiments.

### ELCA detects small molecule-induced modulation of glycosylation at early timepoints

Having demonstrated a temporal difference in the impact on cell surface versus secreted glycoproteins, we next turned to evaluating the dynamics of commonly used inhibitors of cellular glycosylation, including Benzyl-α-GalNAc (Benz; inhibiting *O*-GalNAc extension) and Kifunensine (Kif; mannosidase I inhibition, causing accumulation of high-mannose *N*-glycans). To this end, we analyzed cell surface glycosylation by flow cytometry and secreted glycoproteins by ELCA in parallel from the same experimental samples (Fig. 3A).

**Figure 3.**
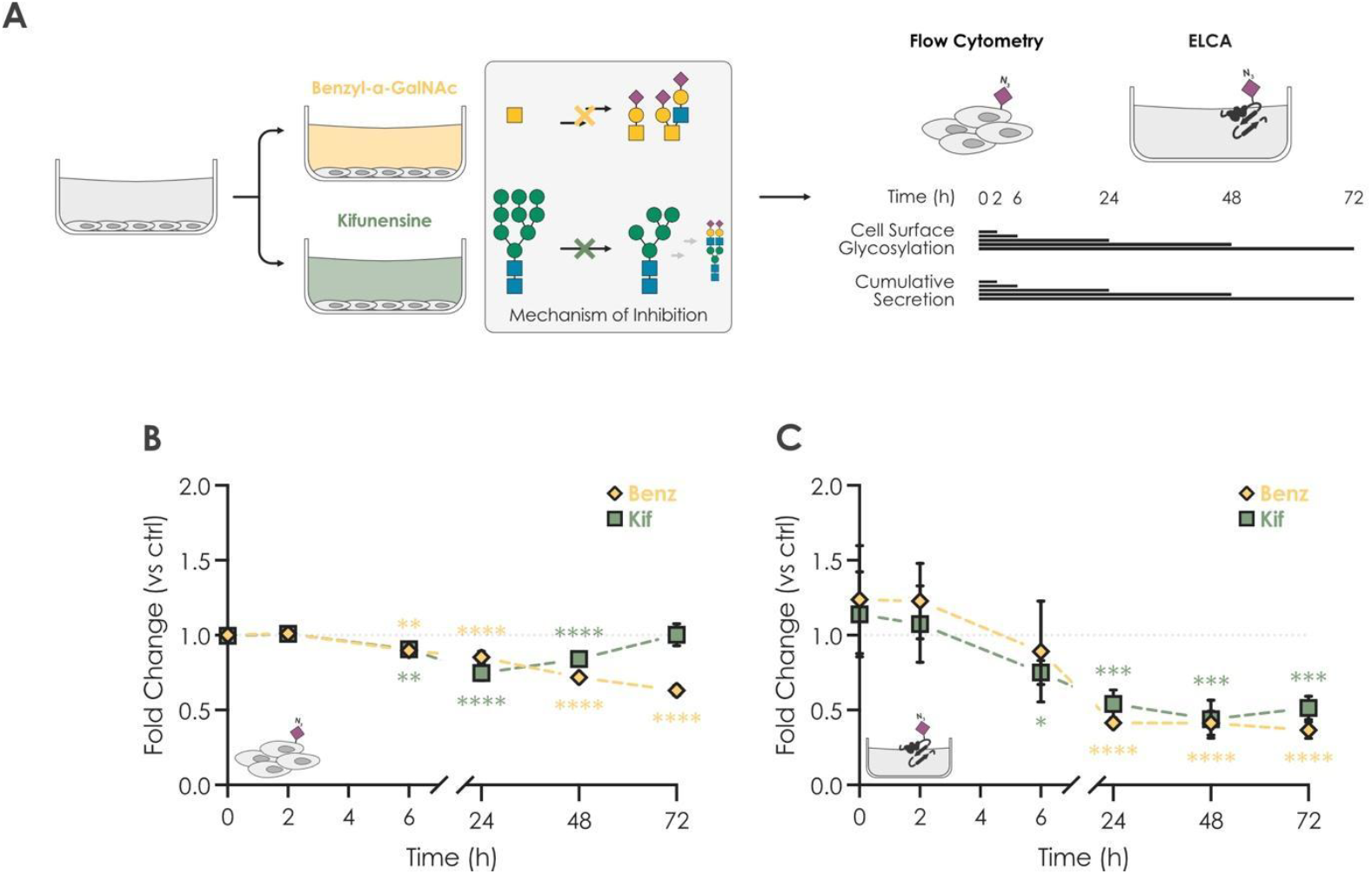
ELCA analysis of culture supernatant is superior to cell surface flow cytometry for detecting small molecule-induced modulation of glycosylation. **A)** Overview of the experimental setup. A375 cells were metabolically labeled with 50 µM Ac_4_ManNAz for 72 h, followed by re-seeding in fresh serum-containing medium supplemented with common glycosylation inhibitors (1 mM Benzyl-α-GalNAc, Benz, and 10 µM Kifunensine, Kif). At the indicated time points, supernatants were collected and stored at −20 °C until ELCA analysis. Cells were washed once with Opti-MEM, then incubated with 10 µM DBCO-biotin for 1 h at 37 °C and subsequently analyzed by flow cytometry. **B)** Flow cytometry analysis of biotinylated cell surface glycoproteins using fluorescein-conjugated streptavidin for detection (n = 4). **C)** ELCA measurements of serum-containing supernatants using high-sensitivity luminescent detection (n = 4), reflecting cumulative protein secretion for the duration of the treatment. All assays were conducted in A375 cells. Ctrl: untreated control. Statistical significance was assessed using two-way ANOVA with Tukey's multiple comparisons test (<2 conditions, 2 variables). *p < 0.05, **p < 0.01, ***p < 0.001, ****p < 0.0001.

Comparing the relative effects on cell surface and secreted glycoproteins in a time-course experiment, we found that treatment with Benzyl-α-GalNAc induced a gradual, time-dependent change in cell surface glycosylation. A significant but modest decrease was detectable after 6 h of treatment, reaching a maximal reduction of approximately 40% after 72 h (Fig. 3B). In contrast, ELCA analysis of supernatant samples revealed more rapid and pronounced effects, with a sustained 60-75% reduction in signal observed throughout the 24-72 h time points (Fig. 3C).

Analysis of cell surface glycosylation by flow cytometry showed that Kifunensine treatment resulted in a maximal signal reduction of approximately 25% after 24 h relative to the untreated control (Fig. 3B), with signal levels returning to baseline by 72 h. In contrast, ELCA measurements of supernatant samples revealed a substantially greater effect, with signal reductions of up to ~60% that persisted throughout the treatment period (Fig. 3C). We thus caution researchers using these inhibitors to study membrane-associated glycosylation, as sufficient timescales may be required to allow for equilibration in such experiments (e.g., Benzyl-α-GalNAc), while, at the same time, transient effects may be overlooked if sampling is performed outside the optimal detection window (e.g., Kifunensine).

### Quantifying secreted glycoproteins from THP-1-derived macrophages

To benchmark ELCA in a physiologically relevant model, we quantified secreted glycoproteins from THP-1-derived macrophages under resting state and during differentiation induced by polarizing treatments. Specifically, we compared naïve M0 macrophage-like cells (PMA-differentiated THP-1), classically activated (M1) macrophages induced by LPS, and alternatively activated (M2) macrophages induced by IL-4/IL-13 (Fig. 4A). These conditions are known to generate cell populations with distinct cytokine profiles and downstream effector functions.

**Figure 4.**
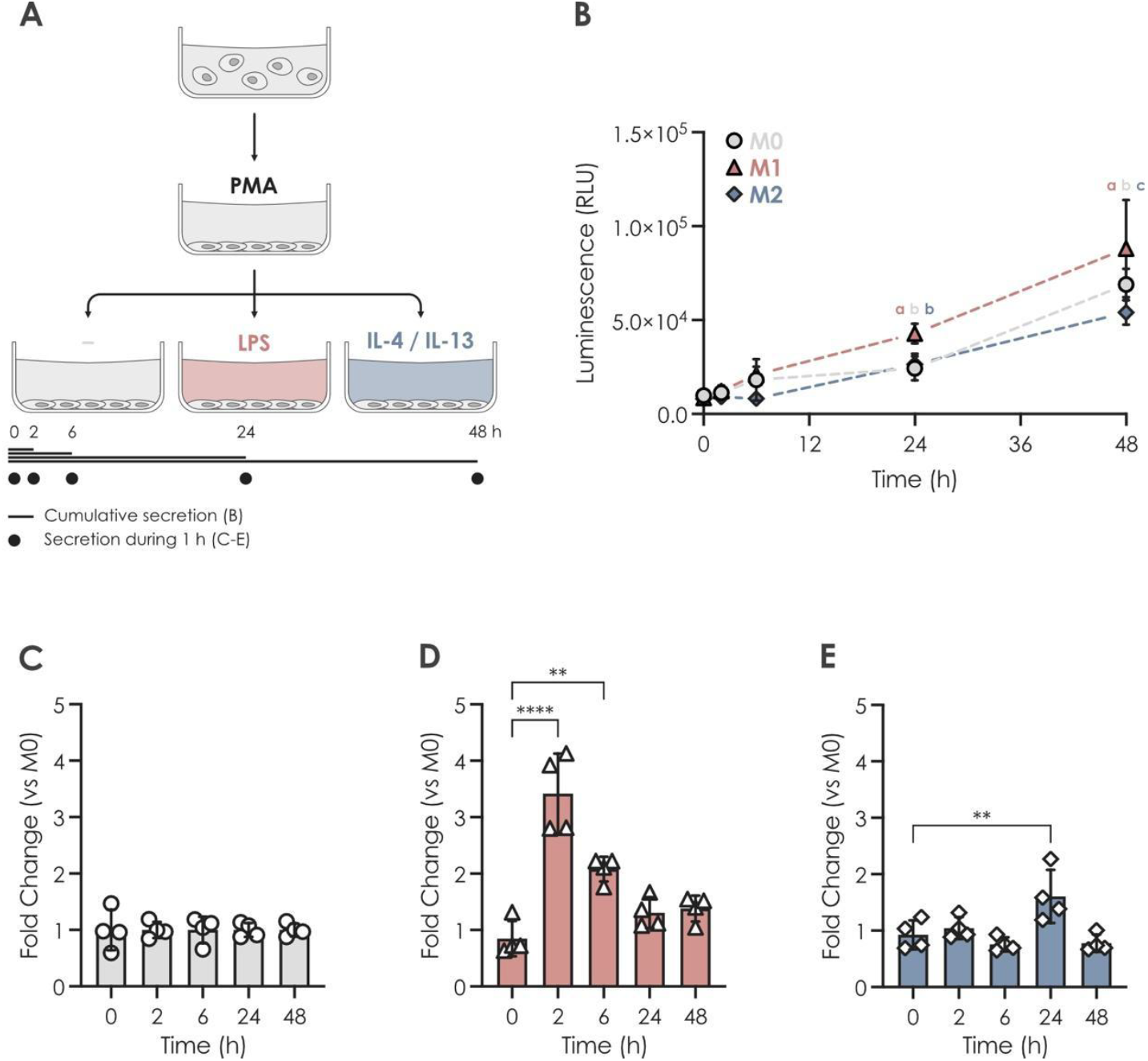
ELCA detects differential secretion profiles during THP-1-derived macrophage polarization. **A)** Overview of the experimental setup. THP-1 cells were differentiated into naïve macrophage-like (M0) cells and metabolically labeled (20 nM PMA, 50 µM Ac_4_ManNAz for 72 h), followed by 24 hours of rest and subsequent activation with LPS (100 ng/mL, M1) or IL-4 and IL-13 (20 ng/mL each, M2). At the indicated time points, serum-containing supernatants were collected and stored at −20 °C until ELCA analysis. Cells were then incubated in fresh serum-free Opti-MEM for 1 h, after which supernatants containing proteins secreted during this period were collected and stored at −20 °C for ELCA analysis. **B)** ELCA measurements of serum-containing supernatants using high-sensitivity luminescent detection (n = 4), reflecting cumulative protein secretion for the duration of the treatment. At each timepoint, letters indicate statistical groupings. **C-E)** ELCA measurements of serum-free supernatants using high-sensitivity luminescent detection (n = 4), reflecting the abundance of proteins secreted during the 1 h incubation period. All assays were conducted in THP-1 cells. Statistical significance was assessed using one-way ANOVA with Dunnett’s multiple comparisons test (>2 conditions, 1 variable) or two-way ANOVA with Tukey’s multiple comparisons test (>2 conditions, 2 variables). **p < 0.01, ****p < 0.0001.

Measurements of cumulative secretion at the indicated time points throughout the experiment revealed a significantly higher abundance of secreted glycoproteins in LPS-activated M1 macrophages from 24 h onwards (Fig. 4B). At each time point, we also measured secretion over a 1 h period to assess secretion rates across conditions (Fig. 4C-E). In strong agreement with previous reports on human macrophage activation^27^, we observed a strong, early response to M1-polarizing treatment (Fig. 4D), whereas M2-polarizing treatment resulted in a more gradual and delayed response (Fig. 4E).

## Discussion

ELCA offers a flexible approach for detecting and quantifying secreted glycoproteins. Cells can be monitored under physiology-mimicking growth conditions, i.e., including the presence of high-abundance FBS-derived (glyco)proteins, while maintaining sufficient sensitivity to detect rapid, small-molecule-induced changes in glycosylation and/or glycoprotein abundance.

Detection sensitivity can be further enhanced by using serum-free conditions in experiments where this is tolerated. In addition, cells may first be cultured under standard conditions, preserving viability during long-term treatments, to measure cumulative secretion over the course of the experiment. Then, at the experimental endpoint, secretion rates can be estimated by quantifying proteins secreted into fresh serum-free medium during a defined short-term interval, such as 1 h.

Bioorthogonal metabolic labeling with Ac_4_ManNAz enables the detection and quantification of global sialoglycoproteins. By using alternative azide-functionalized monosaccharide precursors, the method can be adapted to quantify specific glycan classes, as demonstrated here with Ac_4_GalNAz for *O*-GalNAc glycoproteins. Although additional optimization and validation would be required, the approach should also be readily adaptable to an ever-growing plethora of selective bioorthogonal probes, including azide-functionalized fucose analogs for the detection of fucosylated glycans^28^ or 1,3-Pr_2_-6-OTs GlcNAlk for hybrid *N*-glycan structures^29^ (using azide-biotin for detection in the latter case).

Analogous to how exoglycosidase treatment is being used to endow mass spectrometry with higher structural resolution^30^, ELCA could potentially be further customized by pre-treating samples with, e.g., α2-3 linkage-specific sialidases prior to the click reaction to only quantify α2-6 linked sialylation, if needed for a biological question.

While alternative methods can provide detailed structural insights into glycans, ELCA is well suited for initial screening and high-throughput experimental designs involving many replicates and/or conditions. We also envision ELCA to find usage in screening new glycosylation inhibitors that are needed to advance glycobiology. With an approximate cost of 1-2 EUR per sample (assayed in technical duplicates in a 96-well plate format) for standard colorimetric or high-sensitivity luminescent detection, respectively, ELCA is orders of magnitude less expensive than dedicated glycomics or glycoproteomics approaches. At the same time, it remains highly accessible, as it does not require specialized equipment.

As a limitation of our method, we caution that ELCA, by design, does not report on lipid glycosylation, which is only found on the plasma membrane and is not secreted. Further, it is known that the usage of azide-containing sialic acid may shift the distribution of sialyltransferase activity in subtle ways^31^. As standard ELCA merely reports on global sialylation levels, regardless of linkage, we deem this of lesser concern for most applications.

## Methods

### Mammalian cell culture

A375 cells were cultured in Dulbecco’s Modified Eagle Medium (DMEM) with high glucose, 4.0 mM L-glutamine, and sodium pyruvate (SH30243.01, Cytiva). THP-1 cells (TIB-202, ATCC) were cultured in Roswell Park Memorial Institute (RPMI) 1640 medium (ATCC modification; A1049101, Gibco). The respective growth media were supplemented with 10% (v/v) heat-inactivated fetal bovine serum (FBS; FBS-HI-12A, Capricorn Scientific), and 1% (v/v) penicillin-streptomycin solution (P4333-100ML, Sigma-Aldrich). All cells were maintained at 37 °C in a humidified incubator with 5% CO2.

THP-1 cells were differentiated into macrophage-like cells by treatment with 20 nM phorbol-12-myristate-13-acetate (PMA; P8139-1MG, Sigma-Aldrich) for 72 h followed by a 24 h rest period in fresh medium. Classically activated (M1) macrophages were induced by treatment with 100 ng/mL lipopolysaccharide (LPS; L4391-1MG, Sigma-Aldrich) and alternatively activated (M2) macrophages were induced by treatment with 20 ng/mL interleukin 4 (IL-4; SRP3093-20UG, Sigma-Aldrich) and 20 ng/mL interleukin 13 (IL-13; SRP3274-10UG, Sigma-Aldrich).

Unless stated otherwise, metabolic labeling was performed by growing the cells in the presence of 50 µM Ac_4_ManNAz (CLK-1084-25, Jena Bioscience) for 72 h. Small molecule-mediated modulation of cellular glycosylation was performed by treatments with 1 mM Benzyl-α-GalNAc (HY-129389, MedChemExpress) and 10 µM Kifunensine (HY-19332, MedChemExpress). For all treatments, the untreated control included matched concentrations of vehicle solvent.

### Enzyme-Linked Cycloaddition Assay

A step-by-step protocol is available in Supplementary Methods. Briefly, A375 cells were metabolically labeled with Ac_4_ManNAz prior to seeding in 24-well plates (500 µL per well, 0.25 × 10^6^ cells/mL). Supernatants were collected and stored at −20 °C until further analysis. For ELCA, 50 µL of supernatant was mixed with 50 µL of 0.2 M carbonate–bicarbonate pH 9.4 buffer (28382, Thermo Fisher) and adsorbed onto 96-well MaxiSorp flat-bottom plates (clear or white plates for colorimetric or luminescent detection, respectively) overnight at 4 °C. Blocking was performed using SuperBlock (TBS) Blocking Buffer (37535, Thermo Fisher) supplemented with 10% FBS for 2 h at room temperature. Strain-promoted azide-alkyne cycloaddition was carried out by incubating with 10 µM DBCO-PEG4-Biotin (HY-130809, MedChemExpress) or PC DBCO Biotin (CCT-1120-1, Vector Laboratories) in SuperBlock (TBS) Blocking Buffer for 2 h at room temperature with gentle shaking (500 rpm). HRP labeling was performed by incubation with 2.5 ng/mL streptavidin-HRP in SuperBlock (TBS) Blocking Buffer for 1 h at room temperature with gentle shaking. Detection was carried out using either a colorimetric substrate (TMB Substrate Kit; 34021, Thermo Fisher) or a luminescent substrate (SuperSignal ELISA Femto Substrate; 37075, Thermo Fisher), according to the manufacturer’s instructions. Washing steps consisted of five washes with TBS-T, each 1 min incubation with gentle shaking.

For the detection of specific glycosylated targets, ELCA was combined with a standard sandwich ELISA-based antigen capture approach. Briefly, clear 96-well MaxiSorp flat-bottom plates were coated overnight at 4 °C with 2 µg/mL GM-CSF monoclonal antibody (clone 3092; M500A, Thermo Fisher) and subsequently blocked with 1% BSA for 1 h at room temperature. Collected supernatants were reacted with 10 µM DBCO-PEG4-biotin for 2 h at room temperature. 100 µL of click-labeled supernatant was then added per well, and the plate was incubated for 2 h at room temperature. After washing, the plate was incubated with 2.5 ng/mL streptavidin-HRP for 1 h, followed by colorimetric detection as described above.

### XTT assay

20 000 cells (100 µL of 0.2 × 10^6^ cells/mL) were seeded in each well of a 96-well plate. After incubation for 24 hours under the assayed experimental conditions, 50 µL XTT labeling mixture (5 mL XTT labeling reagent + 0.1 mL electron coupling reagent for one full 96-well plate) was added and the plate incubated for 4 h. Background absorbance measured at the reference wavelength 660 nm was subtracted from specific absorbance of the formazan product measured at 450 nm. The obtained values were normalized between standard culture conditions (DMEM + 10% FBS) and a blank medium control without cells and are reported as metabolic activity (%).

### Flow cytometry detection of azide-tagged cell surface glycoproteins

The cells were washed once in Opti-MEM (31985070, Thermo Fisher) and then incubated with 10 µM DBCO-PEG4-Biotin (HY-130809, MedChemExpress) or PC DBCO Biotin (CCT-1120-1, Vector Laboratories) in Opti-MEM for 1 hour at 37 °C with 5% CO_2_. After labeling, the cells were washed in normal culture medium for 1 hour before harvesting. Detached cells were washed once in PBS and then incubated with 2 µg/mL streptavidin-fluorescein (SA-5001-1, Vector Laboratories) for 30 min at 4 °C. Surface staining was done together with LIVE/DEAD Fixable Far Red Dead Cell Stain Kit (L10120, Thermo Fisher, 1000× dilution). Data were acquired on an Accuri C6 Plus Flow Cytometer (BD) and analyzed using FlowJo 10.10.0 software.

### Quantification and statistical analysis

Comparing two groups was done via two-tailed Welch’s t-tests. Comparing more than two groups was done via one-way ANOVA with Dunnett’s multiple comparison test (1 variable) or via two-way ANOVA with Tukey’s multiple comparisons test (2 variables).

## Acknowledgement

This work was supported by a Branco Weiss Fellowship – Society in Science awarded to D.B.; by the Knut and Alice Wallenberg Foundation; the Jeansson Foundations; the Swedish Foundation for Strategic Research; and the University of Gothenburg, Sweden.

## Contributions

D. B. and J. L. conceptualization; J. L. and J. Y. formal analysis; D. B., J. L., and J. Y. writing–original draft; D. B., J. L., and J. Y. writing–review & editing; J. L. visualization; D. B. supervision; D. B. funding acquisition; D. B., J. L., and J. Y. methodology

