## Supplemental Figures for "Enzyme-Linked Cycloaddition Assay (ELCA) for rapid, ultra-sensitive monitoring of secreted sialoglycoproteins"

### Supplementary Figures

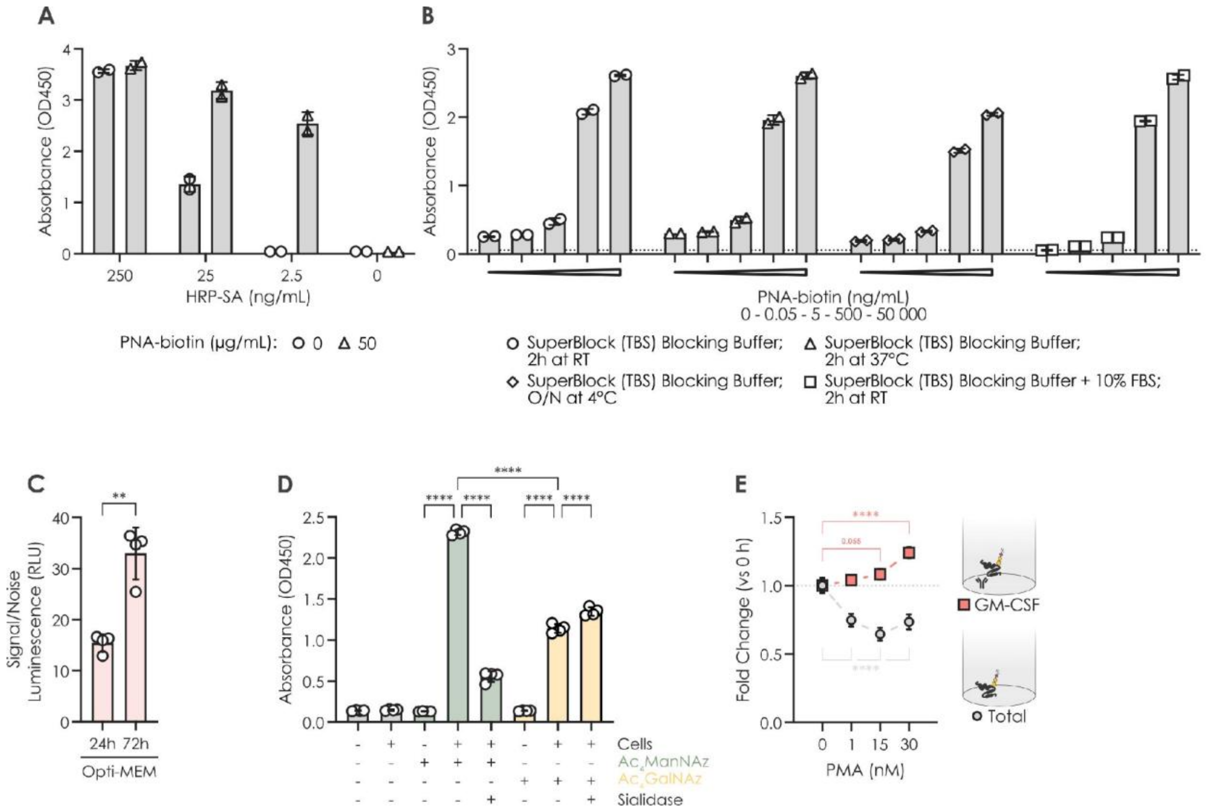


**Figure S1. Quantifying secreted glycoproteins under standard culture conditions. A)** Titration of streptavidin-conjugated horseradish peroxidase (SA-HRP) used for the detection of captured, biotinylated protein (Peanut Agglutinin (PNA), Biotinylated; B-1075-5, Vector Laboratories). The best signal-to-noise ratio is observed at a concentration of 2.5 ng/mL (n = 2). **B)** Comparison of different blocking reagents and protocols. The lowest background signal (indicated by the dotted line) is observed with blocking using SuperBlock Blocking Buffer supplemented with 10% serum for 2 h at room temperature (n = 2). **C)** Signal-to-noise ratio (S/N) of ELCA signal from cells cultured under serum-free conditions at 24 h and 72 h timepoints, measured using high-sensitivity luminescent detection (n = 4). **D)** ELCA signal measured from samples with or without cells, metabolic labeling, and sialidase treatment under serum-free conditions for 24 h, using standard colorimetric detection (n = 4). **E)** ELCA measurements of total glycoprotein secretion (grey) and Sandwich-ELCA measurements of GM-CSF secretion (red) following PMA stimulation in cells cultured under standard conditions for 48 h, using standard colorimetric detection or high-sensitivity luminescent detection, respectively (n = 4). Statistical significance was assessed using a two-tailed Welch's t-test (two conditions), one-way ANOVA with Dunnett's multiple comparisons test (>2 conditions, 1 variable), or two-way ANOVA with Tukey's multiple comparisons test (<2 conditions, 2 variables). **p < 0.01, ****p < 0.0001.

### Supplementary Methods

#### Enzyme-Linked Cycloaddition Assay (step-by-step protocol)

General notes

- Colorimetric readout: Use 96-well clear flat-bottom MaxiSorp plates (439454, Thermo Fisher) and TMB Substrate Kit (34021, Thermo Fisher) for detection.
- Luminescent readout: Use 96-well white flat-bottom Maxisorp plates (436110, Thermo Fisher) and SuperSignal ELISA Femto Substrate (37075, Thermo Fisher) for detection.
- It is recommended to analyze each biological replicate in (at least) technical duplicates.
- Additional materials:
    - Reagent Reservoirs (4870, Corning)
    - Sealing Tape (15036, Thermo Fisher)

Cell culture

1 $Ac_4$ManNAz (CLK-1084-25, Jena Bioscience) metabolic labeling.
   a. 50 µM for 72 h.
   b. May require cell line-dependent optimization for optimal labeling.

2 Prepare samples.
   a. Collect metabolically labeled cells, count, and re-seed 500 µL of $0.25 \times 10^6$ cells/mL cell suspension per well in a 24-well plate with appropriate treatments.
   b. Maintain 50 µM $Ac_4$ManNAz in the culture medium to ensure constant labeling.
   c. Appropriate controls include (i) medium background control; medium containing $Ac_4$ManNAz but no cells, and (ii) DBCO labeling control; supernatant from cells grown/treated in the absence of metabolic labeling.
   d. Optional: Re-seed cells under standard culture conditions or in serum-free medium.
   e. Harvest supernatants after centrifugation to pellet any floating cells and store at −20 °C until analysis.

ELCA measurement

1. Plate adsorption
   a. Add 50 µL 0.2 M carbonate–bicarbonate pH 9.4 (28382, Thermo Fisher) binding buffer per well.
   b. Add 50 µL supernatant sample per well.
   c. Incubate overnight at 4 °C.
2. Wash
   a. 1× with 300 µL 1× TBS-T (28360, Thermo Fisher) per well.
3. Block
   a. Add 300 µL SuperBlock (TBS) Blocking Buffer (37535, Thermo Fisher) supplemented with 10% serum (FBS-HI-12A, Nordic Biolabs) per well.
   b. Incubate 2 h at room temperature.
4. Wash
   a. 1× with 300 µL 1× TBS-T per well.

5. SPAAC labeling of captured azido-glycoproteins
    a. Add 100 µL 10 µM DBCO-Biotin (HY-130809, MedChemExpress) in SuperBlock (TBS) Blocking Buffer per well.
    b. Incubate 2 h at room temperature with gentle shaking (~500 rpm).
6. Wash
    a. 5× with 300 µL 1× TBS-T per well. Incubate 1 min with gentle shaking between each wash.
7. HRP labeling of biotinylated proteins.
    a. Add 100 µL 2.5 ng/ml SA-HRP (N100, Thermo Fisher) in SuperBlock (TBS) Blocking Buffer per well.
    b. Incubate 1 h at room temperature with gentle shaking in the dark.
8. Wash
    a. 5× with 300 µL 1× TBS-T per well. Incubate 1 min with gentle shaking between each wash.
9. Detection
    a. Colorimetric, e.g., TMB Substrate Kit:
        i. Immediately before use, mix equal volumes of the TMB Solution and the Peroxide Solution.
        ii. Add 100 µL of the TMB substrate solution to each well.
        iii. Incubate at room temperature for 30 minutes or until the desired color develops. Optional: Incubate at 37 °C for faster development.
        iv. Stop reaction by adding 100 µL 0.5 M sulfuric acid per well.
        v. Measure the absorbance of each well at 450 nm.
    b. Luminescent, e.g., SuperSignal ELISA Femto Substrate:
        i. Immediately before use, mix equal volumes of the SuperSignal ELISA Femto Luminol/Enhancer and SuperSignal ELISA Femto Stable Peroxide solutions.
        ii. Add 100 µL Working Solution to each well.
        iii. Incubate 1 min with gentle shaking.
        iv. Immediately measure luminescence (~425 nm peak emission, for best sensitivity measure total light output).
